# The limitation of lipidation: conversion of semaglutide from once-weekly to once-monthly dosing

**DOI:** 10.1101/2024.08.10.607458

**Authors:** Eric L. Schneider, John A. Hangasky, Rocio del Valle Fernandez, Gary W. Ashley, Daniel V. Santi

## Abstract

The objective of this work was to develop a long-acting form of the lipidated peptide semaglutide that can be administered to humans once-monthly. Semaglutide was attached to 50 μ diameter hydrogel microspheres by a cleavable linker with an expected in vivo release half-life of about one-month. After a single subcutaneous dose, the pharmacokinetic parameters of released semaglutide were determined in normal mice and the bodyweight loss was determined in diet induced obese mice. The results were used to simulate the pharmacokinetics of semaglutide released from the microspheres in humans.Semaglutide tethered to microspheres by a cleavable linker could be completely released with an in vitro half-life of ∼55 days at pH 7.4. The in vivo half-life of released semaglutide was ∼30 days, and a single dose in diet-induced obese mice resulted in a lean-sparing body weight loss of 20% over 1 month, statistically the same as semaglutide dosed twice daily. Simulations indicated the microsphere-semaglutide would permit once-monthly administration in humans. The microsphere-semaglutide conjugate described here should be suitable for once-monthly dosing in humans, and the same approach should enable conversion of other lipidated peptides from once-weekly to once-monthly administration.

## Introduction

GLP-1 receptor agonists (RA) are mainstays of treatments for T2D and obesity, and are potential therapies for MASH and age-related diseases such as as Parkinson’s and Alzheimer’s (1, 2). Most GLP-1RAs consist of short-lived peptides modified with a fatty acid to create longer-acting “lipidated” peptides (3, 4). The fatty acid reversibly binds to serum albumin, and converts the parent peptide’s half-life from ∼one hour to ∼one week by piggybacking on albumin (4). Notably, Novo Nordisk’s tour de force optimization of peptide half-life extension by lipidation alone has likely achieved a practical upper limit.

It has been reported that persistence in anti-obesity drug use is low (5, 6). Among drugs studied, subcutaneously administered semaglutide showed the highest 1-year persistence, yet only 40% were persistent with the medication (5). A proven method to increase adherence to injectable drugs is by reducing the dosing frequency (7), so it seems desirable and important to increase the half-life of anti-obesity peptides to longer than one week. GLP-1RAs with extended half-lives would also address an unmet need in diseases and patient populations that would benefit from monthly or longer dosing that coincides with doctor visits, such as Parkinson’s and Alzheimer’s.

A yet untried approach to decrease the dosing frequency of anti-obesity peptides is to “override” the half-life limit of lipidated peptides with an alternative half-life extension technology. Thus far, the one-week half-life barrier has not been overcome by PEGylation, polymer encapsulation or Fc fusion (8). However, AMG133 – an anti-GIP mAb conjugated to a GLP-1 agonist – has a 2-week half-life that allows once-monthly dosing (9). Still, there are few technologies that can achieve dosing frequencies of one month or greater.

In our approach to half-life extension, a macromolecular prodrug serves as a subcutaneous depot that slowly releases its drug over a designated, pre-programmed period. Here, a drug is covalently tethered to a long-lived carrier – 50 μ hydrogel microspheres (MS) – by a cleavable linker that dictates the half-life (10, 11); the prodrug is administered subcutaneously and the depot slowly releases the drug by a base-catalyzed β-elimination over a pre-determined period. We believe this is the only technology that can reliably achieve in vivo half-lives of one month or longer for peptides (12, 13). Indeed, the approach has already produced an exenatide GLP-1 agonist with a half-life of one month and with drug remaining above therapeutic levels for up to three months (12). Here, we propose that the very same approach could be used to convert lipidated peptides – semaglutide and others – from once-weekly to once-monthly administration.

In the present work, we attached semaglutide to our MSs by a linker with an expected in vivo cleavage half-life of ∼one month. We determined the pharmacokinetic parameters and weight-loss effects of a single dose in diet induced obese (DIO) mice. Finally, we simulated the pharmacokinetics of our once-monthly semaglutide in humans.

## Methods

### Preparation of N_3_-linker-semaglutide

Semaglutide-Na (43 mg, 10.5 μmol) and N,N-diisopropylethylamine (9 mL, 52 μmol) in 1 mL 9:1 DMF/H_2_O was treated with 5-azido-3,3-dimethyl-1-(methylsulfonyl)-2-pentyl succinimidyl carbonate (14), N_3_-linker(MeSO_2_-)-HSI (11.8 mg, 31.4 μmol). After 3 h, HPLC indicated acylation on the Nα and imidazole of His (30% Nα, 9% imidazole, 59% diacylated). A 0.5 M solution of NH_2_OH-HCl, pH 7 (200 μL), was added and after 16 h HPLC indicated 88% Nα-acylated N_3_-linker(MeSO_2_-)-semaglutide. Preparative HPLC (Phenomenex Jupiter 5 μm 300Å C_18_ 250 × 21.4 mm column, 20-to 100% MeCN in H_2_O + 0.05% TFA) gave purified N_3_-linker(MeSO_2_-)-semaglutide (36 mg, 8.2 mmol, 78%). One peak by HPLC (280 nm); MS [M+3H]^3+^ 1458.4379 (calc. 1458.3979).

### Preparation of MS∼semaglutide

Exemplary, N_3_-linker(MeSO_2_-)-semaglutide (31 mg, 7.1 µmol) in 500 µL of DMF was added to BCN-microspheres (700 μL, 5.9 μM BCN) (13) in MeOH and rotated at 225 rpm for 42 hours, 37 °C. Unreacted N_3_-linker(MeSO_2_-)-semaglutide was removed by washing with 4 mL of DMF, then buffer (10 mM NaOAc, 143 mM NaCl, 0.05% Tween 20, 10 mM Met, pH 5.0), and then exchanged into buffer containing 1.2% of 40k hyaluronic acid. To quantify loading, measured aliquots of slurry were dissolved in 50 mM NaOH, neutralized with 125 mM HEPES, pH 7.4, then assayed for peptide using ∈_280_ = 6790 M^-1^cm^-1^.

### Pharmacokinetics in C57BL/6 mice

Male ∼8 week-old C57BL/6 mice (6/group), were dosed SC in the flank with 50 μL MS∼semaglutide to deliver 400 and 2,000 nmol/kg. Blood samples were drawn from the tail vein at various times to give 3 replicates per timepoint, and processed to plasma with K_2_EDTA/protease inhibitors; semaglutide was quantified by ELISA (BMA Biomedicals #S-1530).

### MS-semaglutide in DIO mice

Male ∼22 week-old C57BL/6 DIO mice (3/group; average weight 47 g), were dosed SC with 50 μL of MS-semaglutide to deliver 200, 660 and 2000 nmol/kg. Body weights, food intake and glucose levels were measured at intervals over 28 d. At study termination, DEXA scan was used to determine the fat and lean mass.

## Results

Semaglutide was carbamylated at the N-terminus with N_3_-linker-HSI to give N_3_-linker-semaglutide in ∼80% yield. The N_3_-linker-semaglutide was then coupled to MS-BCN by SPAAC to give conjugates with 2- and 4 μmol semaglutide/mL MS. Treatment of MS∼semaglutide conjugates under accelerated release conditions at pH 9.4, 37 °C, released semaglutide with a t_1/2_ of 13.3 ±0.5 h, or 1,330 h at pH 7.4.

Concentration vs time plots after single injections of 400- and 2,000 nmol/kg MS∼semaglutide showed dose linearity and a t_1/2_ ∼26 days (**Fig. 2**). The C_max_ and AUC_inf_ dose normalized to 1 nmol/kg were 1.6 nM and 1320 nM*h, respectively. The dose-normalized steady-state AUC_0-28d_ QMo MS∼semaglutide is 1300 nM•h, compared to a reported steady-state AUC_0-30d_ of 1500 nM•h for semaglutide (15), giving a relative bioavailability of 87%. **Fig. 3A** compares the weight loss effects of BID semaglutide and varying single doses of MS∼semaglutide in DIO mice. As shown, BID semaglutide at 10 nmol/kg over 1 Mo – a ∼600 nmol/kg cumulative dose – or single doses of 660- and 2,000 nmol/kg MS∼semaglutide gave ∼20% weight loss, and differences were insignificant. DEXA scan measurements at the end of the experiments showed loss of fat mass rather than lean mass (**Fig. 3B**). As expected (15), both BID semaglutide and single dose MS∼semaglutide lowered blood glucose levels in a dose-dependent manner and suppressed appetite/food intake.

**Figure 1.**
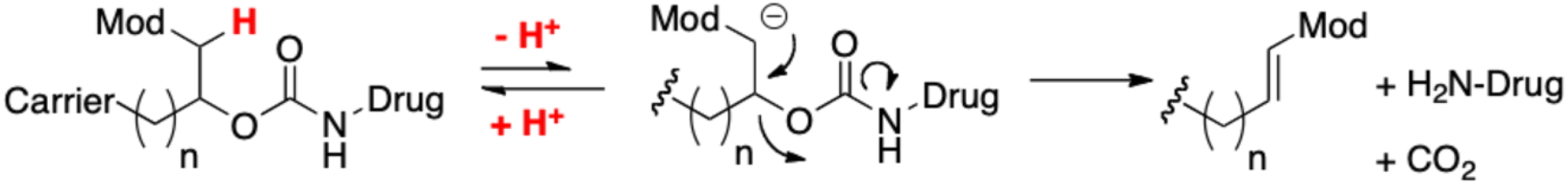
General approach to half-life extension.

**Figure 2.**
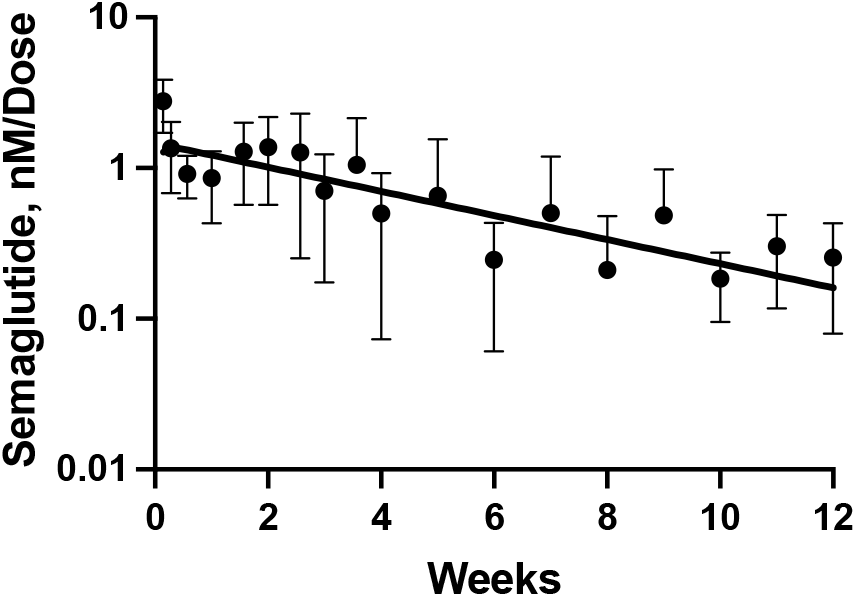
Dose-normalized concentration vs time plot of semaglutide released from MS∼semaglutide in the mouse. Individual data points are dose-normalized mean ± SD (n= 3-to 12) from single doses of 400- and 2000 nmol/kg MS∼semaglutide.

**Figure 3.**
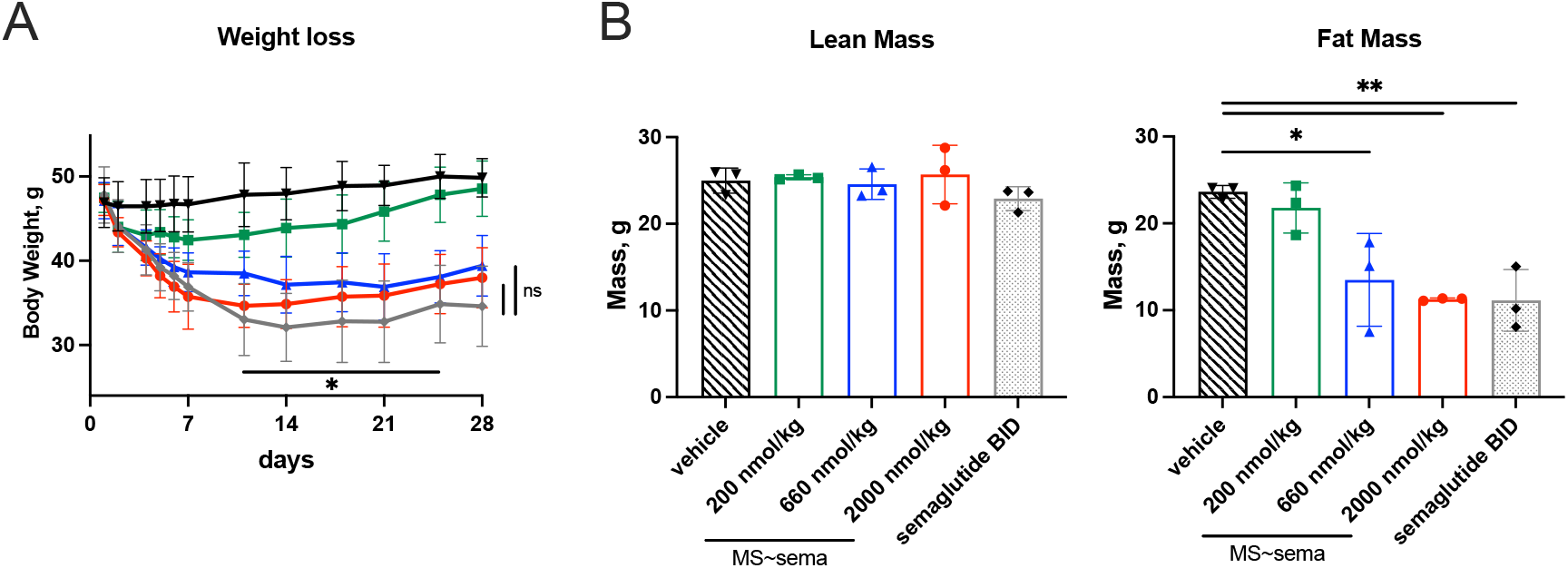
Semaglutide treatment of obese mice. DIO mice 25 wks old and ∼50 g each were treated with vehicle 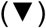, 10 μmol/kg BID semaglutide (600 nmol/kg/mo) 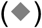 and single doses of MS∼semaglutide 200-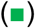, 660-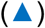 or 2000 nmol/kg 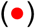 (n=3/group). A) Body weight vs time of DIO mice. Data points are mean ± SD. There is no significant difference between BID semaglutide, or single-dose 660- and 200 MS∼semaglutide; mixed-effects model and Tukey’s post-hoc. B) Composition of weight loss after 28 d treatment. The mice in panel A were analyzed by DEXA scan after 28 d treatment. Data are mean ± SD, and analyzed using 1-way ANOVA and Tukey’s post-hoc. \**p*<0.05, \*\**p*<0.01 vs vehicle.

The dose of MS∼semaglutide at steady state in humans can be estimated from the single dose in mice as follows. First, the effective dose of semaglutide in humans (6 nmol/kg/wk) is 23-fold lower than in mice (140 nmol/kg/wk) (16, 17). From the t_1/2_ of ∼1 Mo, it would take 2.3 Mo for MS∼semaglutide to reach steady state; so, the equi-effective steady state dose of MS∼semaglutide in DIO mice would be ∼2.3-fold lower than the single dose. Therefore, the steady state dose in humans should be 2.3 × 23 or ∼50-fold lower than in mice. We estimate the effective 0.66-to 2.0 μmol/kg/month MS∼semaglutide in mouse is equivalent to 0.013-to 0.04 μmol/kg/month in humans. Hence, for a 100 kg human the effective dose should be ∼0.33-to 1 mL of the 4 μmol/mL preparation of MS∼semaglutide.

## Discussion

We have described an approach whereby a lipidated peptide with a half-life of ∼one week can be converted to a longer-acting peptide with a half-life of ∼one month. The lipidated peptide is tethered to a microsphere depot by a releasable linker that over-rides lipidation and dictates the half-life of the released peptide (12). Here, we demonstrated conversion of semaglutide – which has a half-life of 160 hours – to a prodrug in which the released semaglutide has a half-life of ∼630 h, suitable for QMo administration. And, since the half-life obtained with the technology is species-independent, it should directly translate from mouse to man (18). The exposure and bioavailability of a single dose of MS∼semaglutide vs QD semaglutide over one month are nearly identical, indicating stability and efficient release of semaglutide from MS∼semaglutide in the SC space. Thus, it expected that the pharmacodynamics will likewise be similarly long-acting. Indeed, a single dose of MS∼semaglutide to DIO mice gave a 20% lean-sparing weight loss over one month which is the same as that achieved by a similar cumulative dose of semaglutide given BID over the same period (16). Likewise, MS∼semaglutide caused the appropriate decrease in food intake and glucose (15). Overall, the pharmacokinetic data and protracted period of weight loss show that the linker of MS∼semaglutide – not the lipid – controls the in vivo duration of the drug. Using data obtained in the DIO mouse studies and the effective doses of semaglutide in mouse and man, we estimate the human dose of QMo MS∼semaglutide to comprise of ∼5-to 15 mg semaglutide, which is contained in 0.3-to 1 mL of our 4 μmol/mL MS∼semaglutide preparation. We posit that the very same approach could be used to convert other lipidated peptides of current interest from once-weekly to once-monthly administration. Importantly, the availability of once-monthly anti-obesity agents should mitigate the problem of low persistence, and address an unmet need in patient populations that would benefit from less frequent dosing of a GLP-1 receptor agonist.

## Notes

### Competing Interest Statement

All authors are employees and hold options or stock in ProLynx Inc

